# Evolutionary Divergence of mTOR-mediated Transcriptional Regulation Between *Drosophila melanogaster* and *Drosophila simulans* is modulated by sex and tissue

**DOI:** 10.64898/2026.01.12.699128

**Authors:** Yevgeniy Raynes, John C. Santiago, Faye A. Lemieux, David M. Rand

**Author notes:** **Authors for Correspondence:** Yevgeniy Raynes, Department of Ecology, Evolution, and Organismal Biology, Brown University, Providence, USA,; David Rand, Department of Ecology, Evolution, and Organismal Biology, Brown University, Providence.

## Abstract

Evolutionary divergence in gene regulation is a major source of phenotypic novelty, yet understanding how deeply conserved, pleiotropic signaling pathways evolve under selective constraint while maintaining essential cellular functions remains an important challenge. Here, we investigate the evolutionary divergence in transcriptional regulation mediated by the mechanistic target of rapamycin (mTOR) pathway in two related species, *Drosophila melanogaster* and *Drosophila simulans*. mTOR is a highly conserved central regulator of cellular growth and metabolism, making it well-suited for the study of regulatory evolution under functional constraint. Using a fully factorial RNA-seq design, we quantified transcriptional responses to mTOR inhibition by rapamycin across three tissues (head, thorax, and abdomen) and both sexes. Despite highly conserved tissue-specific expression – reflecting the close phylogenetic relationship between species – mTOR-mediated transcriptional responses showed clear evidence of evolutionary divergence. Divergence varied across tissues and sexes: heads of both sexes and female thoraces and abdomens showed the most rapid gene-level divergence, suggesting stronger directional selection, whereas male thorax and abdomen exhibited comparatively conserved responses, consistent with stabilizing selection. Gene-level divergence patterns were consistent across multiple metrics and generally mirrored pathway-level divergence, except in the female abdomen, which showed relative pathway-level conservation despite extensive gene-level divergence. Species-by-treatment interaction analyses further revealed divergence in core mTOR-regulated biological processes. Together, our results suggest that regulatory modularity may allow even highly conserved signaling pathways to evolve under context-specific selective pressures while maintaining critical functionality.

**Significance statement:** The evolution of gene regulation is increasingly recognized as a major contributor to biological diversification. However, how highly pleiotropic and deeply conserved pathways evolve new regulatory effects without disrupting essential cellular functions remains an open question in evolutionary biology. Using the nutrient-sensing mTOR pathway as a model, we show that gene regulation mediated by this core, selectively-constrained signaling network has diverged between two closely related fruit fly species in a tissue- and sex-specific manner. Our findings suggest that even pleiotropic pathways like mTOR can diverge over short evolutionary timescales, and that regulatory modularity may facilitate the evolution of novel transcriptional effects in specific contexts while preserving essential functions.

## Introduction

Evolutionary innovation can be driven by changes in protein-coding sequences that produce new or modified genes, or by regulatory changes that alter the timing, location, or magnitude of gene expression. Regulatory divergence is now recognized as a key mechanism of phenotypic diversification, adaptation, and speciation (King and Wilson 1975; Wray 2007; Tirosh, et al. 2009; Romero, et al. 2012; Signor and Nuzhdin 2018). Indeed, new regulatory patterns, arising from alterations in cis-regulatory elements, trans-acting factors, or chromatin architecture, may generate novel phenotypes while preserving conserved protein machinery (Carroll 2008; Wittkopp and Kalay 2011). Such flexibility is particularly important for pleiotropic pathways, in which changes in coding sequence may be constrained by stabilizing selection on other functions (Wagner and Zhang 2011; Wittkopp and Kalay 2011; Hughes and Leips 2017). Understanding the evolutionary dynamics of transcriptional regulation, especially within essential signaling pathways under strong selective constraint, is therefore critical for linking regulatory divergence to phenotypic novelty.

Here, we use the mechanistic target of rapamycin (mTOR) pathway as a model system for the evolution of transcriptional regulation. mTOR is a serine/threonine kinase that integrates environmental and intracellular cues – such as nutrient availability, energy status, and growth factors – to regulate cell growth, metabolism, and autophagy (Saxton and Sabatini 2017; Kim and Guan 2019). mTOR functions as part of two major complexes. mTOR complex 1 (mTORC1) regulates the balance between anabolism and catabolism, stimulating protein, lipid, and nucleotide biosynthesis and reducing autophagy (Laplante and Sabatini 2012; Gonzalez and Hall 2017; Saxton and Sabatini 2017). mTOR complex 2 (mTORC2) regulates cytoskeletal organization and promotes cell survival. Dysregulation of mTOR signaling has been implicated in a wide range of human diseases, including cancer, metabolic disorders, and age-related neurodegenerative conditions such as Alzheimer’s and Parkinson’s disease (Sato, et al. 2010; Castets, et al. 2013; Grabiner, et al. 2014; Perluigi, et al. 2015; Hua, et al. 2019; Querfurth and Lee 2021).

mTORC1, in particular, offers several distinct advantages for investigating evolutionary divergence in core regulatory pathways. First, in addition to its well-characterized effects on translation, it has been shown to have wide-ranging gene regulatory effects, modulating expression levels of enzymes in metabolic pathways involved in biogenesis of ribosomes, mitochondria, and lysosomes, as well as nucleotide and lipid synthesis (Duvel, et al. 2010; Laplante and Sabatini 2013; Ben-Sahra and Manning 2017; Saxton and Sabatini 2017). Second, the core architecture of the mTOR signaling network is evolutionarily ancient and has been conserved across plants, fungi, and animals (van Dam, et al. 2011; Maegawa, et al. 2015; Dobrenel, et al. 2016; Gonzalez and Hall 2017; Saxton and Sabatini 2017; Shi, et al. 2018; Brunkard 2020), reflecting mTOR’s critical role in regulating core cellular processes. Indeed, orthologs of key components of mTORC1 (mTOR, raptor, and mLST8) have been identified in yeast (Loewith and Hall 2011), algae (Perez-Perez, et al. 2017), and all sequenced plant species (Shi, et al. 2018). And finally, mTORC1 activity can be selectively inhibited by rapamycin, a small molecule that binds FKBP12 to allosterically suppress mTORC1 (Li, et al. 2014). Rapamycin has been used extensively in functional analyses of mTORC1 (Hardwick, et al. 1999; Bjedov, et al. 2010; Saxton and Sabatini 2017; Dobson, et al. 2018), providing a foundation for interpreting the divergence in downstream transcriptional effects of mTOR signaling in related species.

In this study, we use two closely related fruit fly species, *Drosophila melanogaster* and *Drosophila simulans*, to examine divergence in mTOR-mediated transcriptional regulation by comparing the effects of mTOR inhibition with rapamycin. *D. melanogaster* and *D. simulans* diverged approximately 2 million years ago (Markow 2015) and retain substantial genetic and phenotypic similarity (Davis, et al. 1996b). However, they differ in ecology, physiology, behavior, and life history traits (David, et al. 2004; Gibert, et al. 2004), and have been widely used as a comparative system for studying transcriptional divergence (Graze, et al. 2009; Wayne, et al. 2011; Coolon, et al. 2014; Cridland, et al. 2020). Using RNA sequencing, we characterized transcriptional responses to rapamycin in three tissues of male and female *D. melanogaster* and *D. simulans* to quantify the tempo and mode of evolutionary divergence in mTOR-regulated transcription. This fully factorial design allowed us to assess whether evolutionary divergence in mTOR-mediated gene regulation is broadly uniform across biological contexts, or instead reflects more modular, context-dependent evolutionary patterns. Our analyses reveal that mTOR-mediated gene regulation has diverged in a sex- and tissue-specific manner, with some tissues exhibiting significantly more rapid divergence in mTOR effects than others and, overall, greater divergence in females than in males. We also find evidence of divergence in several core mTOR-regulated pathways, highlighting how even conserved signaling pathways can evolve novel regulatory effects over relatively short evolutionary timescales.

## Results

### Patterns of Transcriptional Variation Across Tissues, Sexes, and Species

Comparative analyses of the transcriptomes showed substantial variability in gene expression (Figure 1, Supplementary Figure S1, the read count table is available as Supplementary Table S1), with tissue identity accounting for the largest proportion of transcriptional variance (PERMANOVA: R^2^ = 0.486, *F* = 65.17, *p* < 0.0001), followed by sex (R^2^ = 0.143, *F* = 38.35, *p* < 0.0001), species (R^2^ = 0.127, *F* = 34.15, *p* < 0.0001), and, to a lesser extent, rapamycin treatment (R^2^ = 0.004, *F* = 1.03, *p* = 0.366). Consistent with prior work on the evolutionary conservation of organ-specific transcriptional programs (Sudmant, et al. 2015), we observed greater differentiation among tissues within a species than between homologous tissues in different species (Figure 1). Notably, the relative impact of sex and species on transcription varied by body part (Supplementary Figure S1). Species accounted for most of the variance in the head (primarily composed of neuronal tissue in both sexes) and thorax (primarily composed of muscle). Meanwhile, the difference between sexes had the greater influence in the abdomen, which comprises a variety of tissues, most notably the sex-specific reproductive organs. Finally, rapamycin had a comparatively modest impact on genome-wide transcription compared to these major biological factors, consistent with its more targeted influence on a single, though highly pleiotropic, signaling pathway (Supplementary Figure S1).

**Figure 1:**
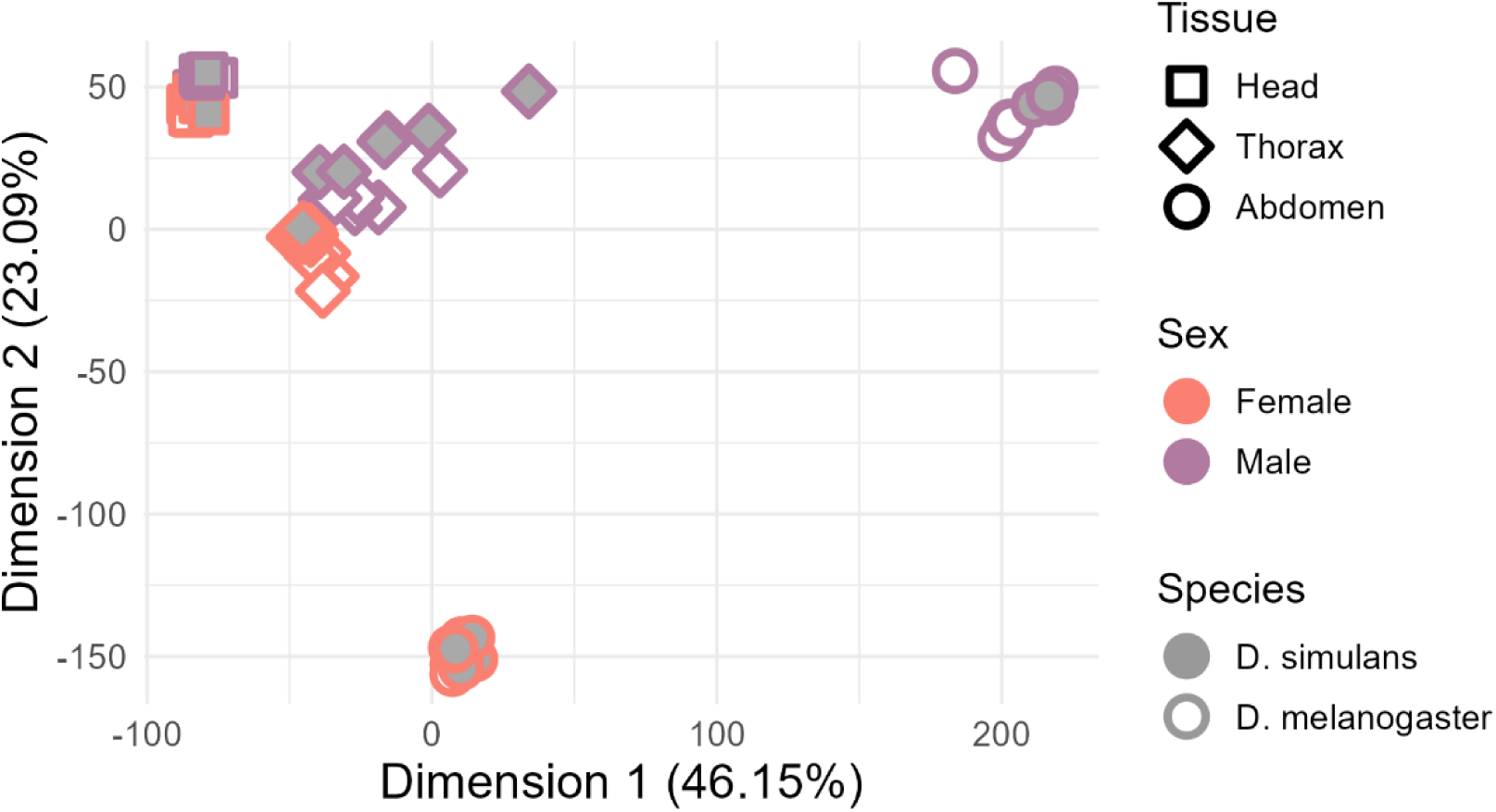
Multidimensional scaling (MDS) analysis of variance-stabilized transcriptome data. MDS reveals substantial variability in gene expression among the three body parts and between the sexes (particularly in the abdomen), and smaller differences between the species and treatments.

Rapamycin effects on expression reveal sex- and tissue-dependent variability in evolutionary divergence of mTOR-mediated transcription.

To quantify divergence in mTOR-regulated transcription, we evaluated the *first-order effects* of rapamycin treatment on transcription in *D. simulans* and *D. melanogaster*. Expression data were partitioned by sex and tissue into six sex-by-tissue combinations, each comprising samples from both species on and off rapamycin. Using DESeq2 (Love, et al. 2014) we identified differentially expressed genes (DEGs) and estimated rapamycin-induced log_2_ fold changes (log_2_FC) in gene expression in each of the sex-by-tissue combinations in *D. melanogaster* and *D. simulans* (Figure 2, Supplementary Figure S2, Supplementary Table S2). To measure divergence in rapamycin response between species, we compared log_2_FC vectors (e.g., values on the x- and y- axes in Figure 2A) using two complementary distance metrics (Glazko and Mushegian 2010): (1) *d_cor_* = 1 – ρ, where ρ is the Spearman’s correlation coefficient, which captures differences in the relative ordering of gene responses, and (2) Euclidean distance (dE), which is sensitive to differences in response magnitude. For both metrics, larger values indicate greater divergence in rapamycin-induced transcriptional responses between species.

**Figure 2:**
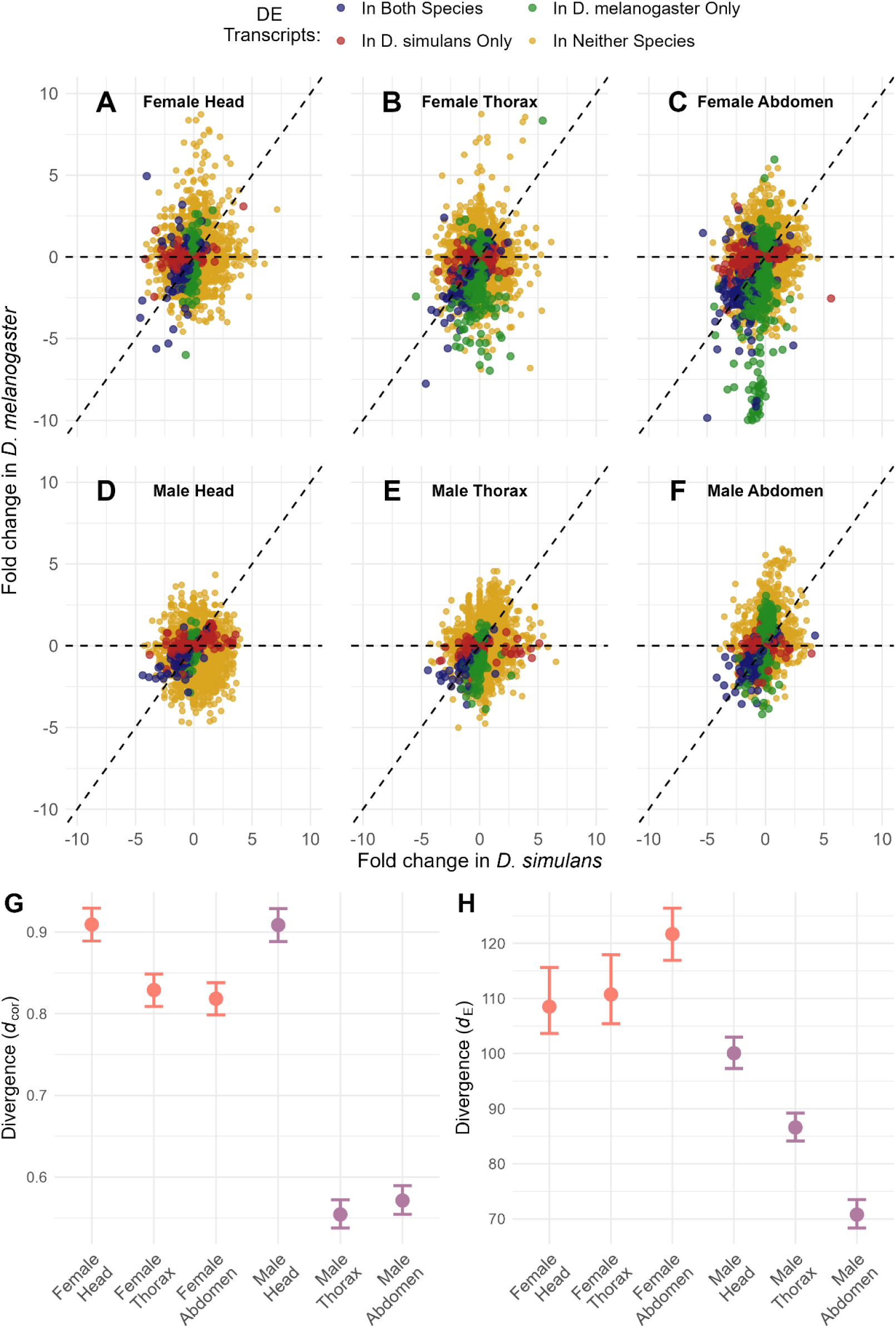
Gene-level divergence in rapamycin transcriptional effects: A-F) log_2_FC in expression in response to rapamycin in *D. melanogaster* and *D. simulans*. Colors indicate whether a transcript is significantly differentially expressed in either, both, or neither species. G) Divergence in the pattern of rapamycin gene response as measured by *d_cor_* (1 – ρ, where ρ is the Spearman’s correlation coefficient) between *D. melanogaster* and *D. simulans* log_2_FC values. H) Divergence in the *magnitude* of response as measured by *d_E_* (Euclidean distance). All panels include all orthologs, regardless of statistical significance or the log_2_FC magnitude. Error bars in panels G and H represent 95% confidence intervals derived from bootstrap resampling (10,000 replicates) with replacement.

Both metrics, calculated across all expressed orthologs shared between species, revealed substantial heterogeneity in transcriptional divergence across tissues and sexes (Figure 2). As measured by *d_cor_* (Figure 2G), heads of both males and females (*d_cor_* ≈ 0.91) exhibited the highest levels of divergence in rapamycin response. Female thorax and abdomen (*d_cor_* ≈ 0.81-0.82) showed intermediate levels of divergence, and male thorax and abdomen were the most conserved (*d_cor_* ≈ 0.55-0.57). Permutation tests (10,000 iterations) showed that all pairwise differences in *d_cor_* were significant (*p* < 0.01), except those between male and female heads (*p* ≈ 0.97), female abdomen and thorax (*p* ≈ 0.45), and between male abdomen and thorax (*p* ≈ 0.19).

As a baseline comparison, we also calculated *d_cor_* divergence in transcriptome-wide expression levels (under control conditions). Consistent with the close phylogenetic relationship between *D. simulans* and *D. melanogaster*, baseline expression divergence was substantially lower than divergence in rapamycin response and showed less variation across tissues or sexes (female heads *d_cor_* ≈ 0.09 ± 0.01, thoraxes *d_cor_* ≈ 0.10 ± 0.01, abdomen *d_cor_* ≈ 0.08 ± 0.01; male heads *d_cor_* ≈ 0.09 ± 0.01, thoraxes *d_cor_* ≈ 0.10 ± 0.01, abdomen *d_cor_*≈ 0.14 ± 0.01; ± 95% CI estimated by bootstrap resampling). Note that baseline expression *d_cor_* and rapamycin-response *d_cor_* are calculated differently (former on absolute expression levels, latter on log_2_FC) and are not directly comparable (baseline *d_cor_* were calculated as a reference for overall transcriptomic divergence).

Divergence quantified by *d_E_*(Figure 2H) was again highest in female tissues (*d_E_* ≈ 108-121), followed by male heads (*d_E_* ≈ 100), and lowest in male thorax (*d_E_* ≈ 81) and abdomen (dE ≈ 71). Permutation tests showed that female abdomen had a significantly higher level of divergence than any other tissue (*p* < 0.001), and that all other pairwise comparisons were significant except between the female head and thorax (*p* ≈ 0.62). We repeated these comparisons using only genes with |log_2_FC| > 0.58 (reflecting an approximately 1.5-fold change in expression) in at least one species, as well as using only DEGs with statistically significant log_2_FC (FDR < 0.05) in at least one species (Supplementary Figure S3). Both approaches revealed broadly similar divergence patterns, with female tissues exhibiting higher levels of divergence than male tissues, and male abdomen and thorax being the most evolutionarily conserved.

We next evaluated overlap between rapamycin-affected DEGs in *D. melanogaster* and *D. simulans* (Figure 3A) using Jaccard dissimilarity (1− *J*) to quantify divergence between the species; *J* is the Jaccard similarity index, calculated as the ratio of shared DEGs (dark blue in Figure 3A) to the total number of unique DEGs identified across both species (i.e., the total number of DEGs in each bar in Figure 3A). Consistent with the divergence patterns observed in log_2_FC distance, female tissues (1−*J* ≈ 0.82 in the head and abdomen; 1−*J* ≈ 0.85 in the thorax) and male heads (1−*J* ≈ 0.83) exhibited the highest dissimilarity in DEG sets, while the male thorax (1−*J* ≈ 0.77) and abdomen (1−*J* ≈ 0.63) were more conserved, even though male abdomen had the most DEGs of all tissues (but also a higher fraction of shared DEGs). Together, log_2_FC distance measures and DEG dissimilarity indicate that the effects of mTOR transcriptional regulation at the level of individual transcripts have diverged at different rates across tissues and sexes. Heads of both sexes appear to be the most labile, followed by female thorax and abdomen, while male thorax and abdomen exhibit comparatively smaller changes.

**Figure 3:**
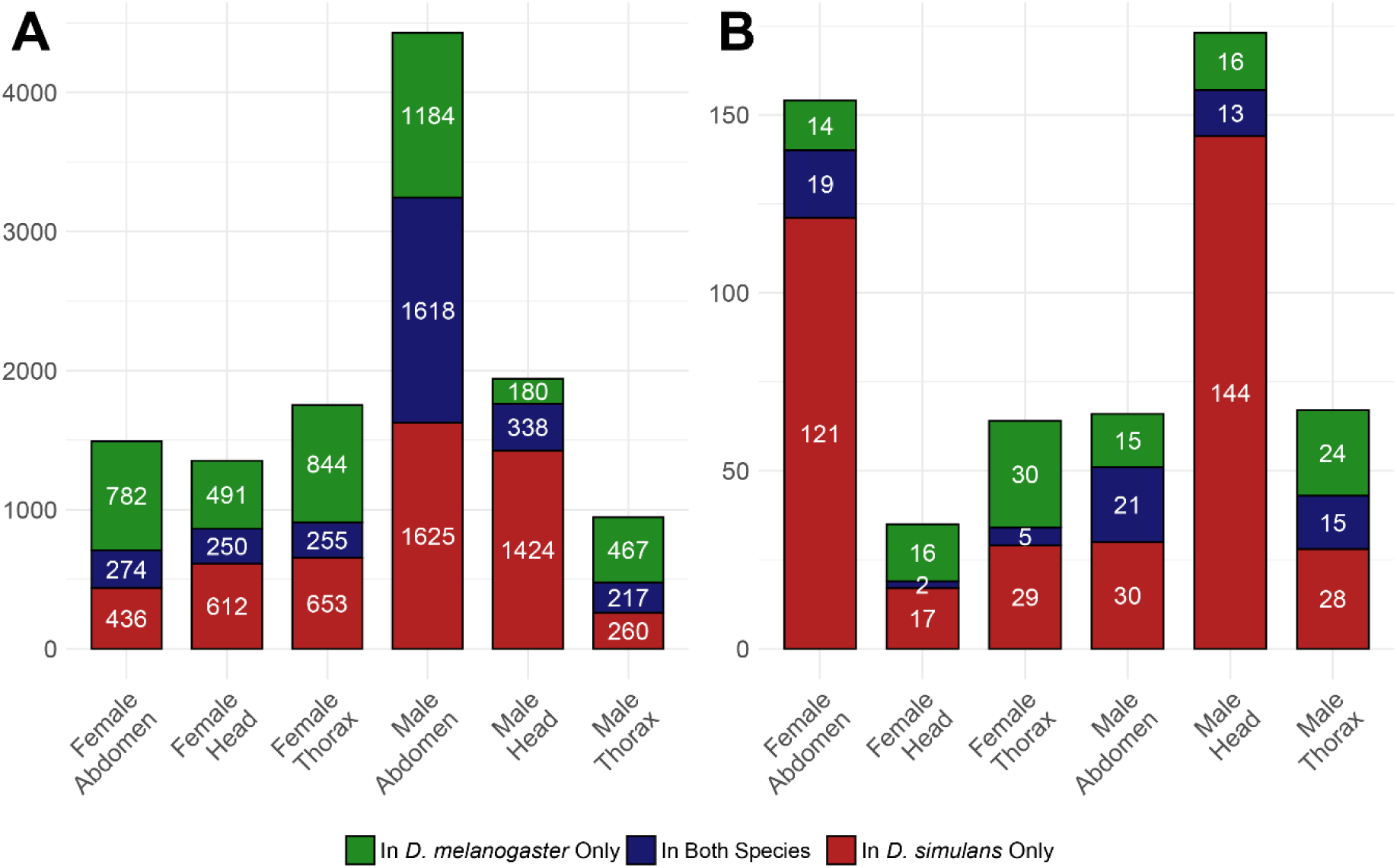
First-order effects of rapamycin treatment. A) Number of significant DEGs shared between species or unique to either *D. melanogaster* or *D. simulans* across sex-by-tissue combinations. B) Number of significantly enriched GO categories identified by gene set enrichment analysis (GSEA) across sex-by-tissue combinations.

### Pathway-level divergence is broadly consistent with gene-level patterns

To assess whether rapamycin-perturbed pathways differ between *D. melanogaster* and *D. simulans* in a sex- or tissue-specific manner, we performed Gene Set Enrichment Analysis (GSEA) of rapamycin effects using Gene Ontology (GO) Biological Process terms (Supplementary Figures S4-S9, Supplementary Table S3). As in the gene-level DEG analyses, we observed both shared and species-specific pathway responses (Figure 3B). We quantified divergence between sets of significantly enriched GO categories in *D. melanogaster* and *D. simulans* (Figure 3B) using Jaccard dissimilarity; *J* was again calculated as the ratio of shared GO categories (dark blue in Figure 3B) to the total number of significant categories identified across both species. Consistent with gene-level divergence patterns, female tissues (1−*J* ≈ 0.94 in the head, 1−*J* ≈ 0.88 in the abdomen, 1−*J* ≈ 0.92 in the thorax) and male heads (1−*J* ≈ 0.92) again exhibited high divergence in enriched GO terms, whereas the male thorax (1−*J* ≈ 0.78) and abdomen (1−*J* ≈ 0.68) were again more conserved.

Notably, the female abdomen and male heads exhibited substantially more significant GO categories in *D. simulans* than in *D. melanogaster*. However, direct comparisons of significantly enriched categories are inherently sensitive to arbitrary significance thresholds. Accordingly, to complement dissimilarity comparisons between significant GO terms, we assessed the distance between normalized enrichment scores (NES) of GO terms in *D. melanogaster* and *D. simulans*. NES provides an estimate of the direction and magnitude of pathway enrichment in response to rapamycin, with negative values indicating pathways enriched among rapamycin-downregulated genes, and positive values indicating pathways enriched among rapamycin-upregulated genes. As before, we used both a correlation-based distance, *d_cor_*, and Euclidean distance, *d_E_* to quantify divergence (Figure 4). When calculated across all GO terms (regardless of statistical significance), both metrics confirmed that heads of both species and the female thorax exhibited the greatest divergence between species (highest *d_cor_* and *d_E_* values), and that male thorax and abdomen were more conserved (especially in Figure 4A), as in previous analyses. Interestingly, the female abdomen exhibited low divergence in NES values despite greater divergence in DEG identities and log_2_FC-based metrics observed previously. These analyses were repeated using only GO terms significantly enriched in at least one species, yielding similar divergence patterns (Supplementary Figure S10).

**Figure 4:**
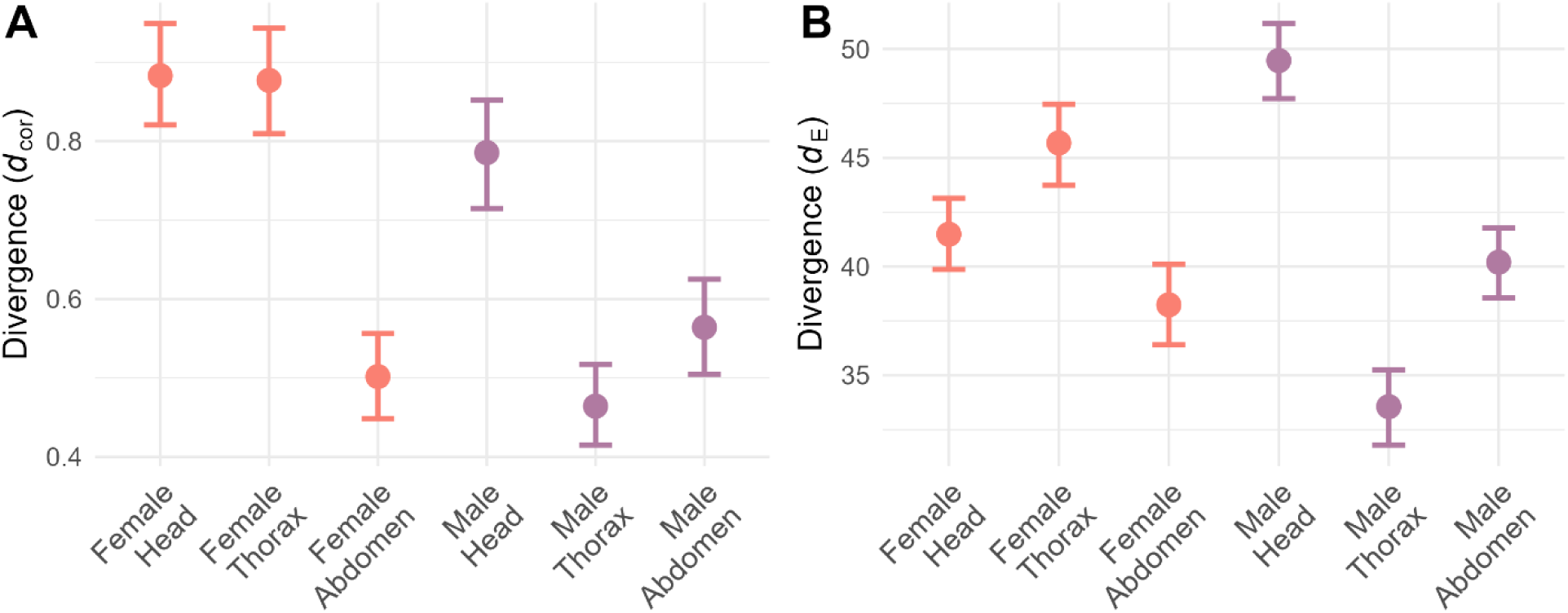
Divergence in rapamycin response at the pathway level. A) *d_cor_* (1 – ρ) between *D. melanogaster* and *D. simulans* NES values for all pathways. B) *d_E_* (Euclidean distance) between *D. melanogaster* and *D. simulans* NES values for all pathways. Error bars show 95% bootstrap confidence intervals (10,000 replicates with replacement).

### Core mTOR-regulated pathways show evidence of species divergence

In addition to quantifying divergence, we aimed to identify pathways exhibiting species-specific transcriptional responses to rapamycin. To do so we evaluated *interaction effects* between rapamycin treatment and species – that is, differences in transcriptional response between *D. simulans* and *D. melanogaster*. Using the same partitioning of the expression data into six sex-by-tissue combinations and a model incorporating both first-order and interaction effects of species and treatment, we identified DEGs with significant interaction terms (Supplementary Figure S11, Supplementary Table S4). To identify species-specific pathway effects of rapamycin, we performed Gene Set Enrichment Analysis (GSEA) on genes ranked by the interaction test statistic (Supplementary Table S5).

GSEA revealed several core mTOR-regulated pathways that were differentially affected by rapamycin in the two species (Figure 5). In female heads, enriched GO terms were dominated by rRNA-related processes, which showed greater suppression in *D. simulans* than *D. melanogaster* (NES < 0). In contrast, pathways related to carbohydrate metabolism and sensory processing were less suppressed by rapamycin in *D. simulans*, as reflected by positive NES values. In male heads, we observed a greater diversity of processes with divergent responses to rapamycin. Core processes involved in protein translation, phosphorylation, and folding, as well as mitochondrial function, were more negatively affected in *D. simulans,* whereas axon guidance, signal transduction, and synaptic transmission were comparatively less suppressed.

**Figure 5:**
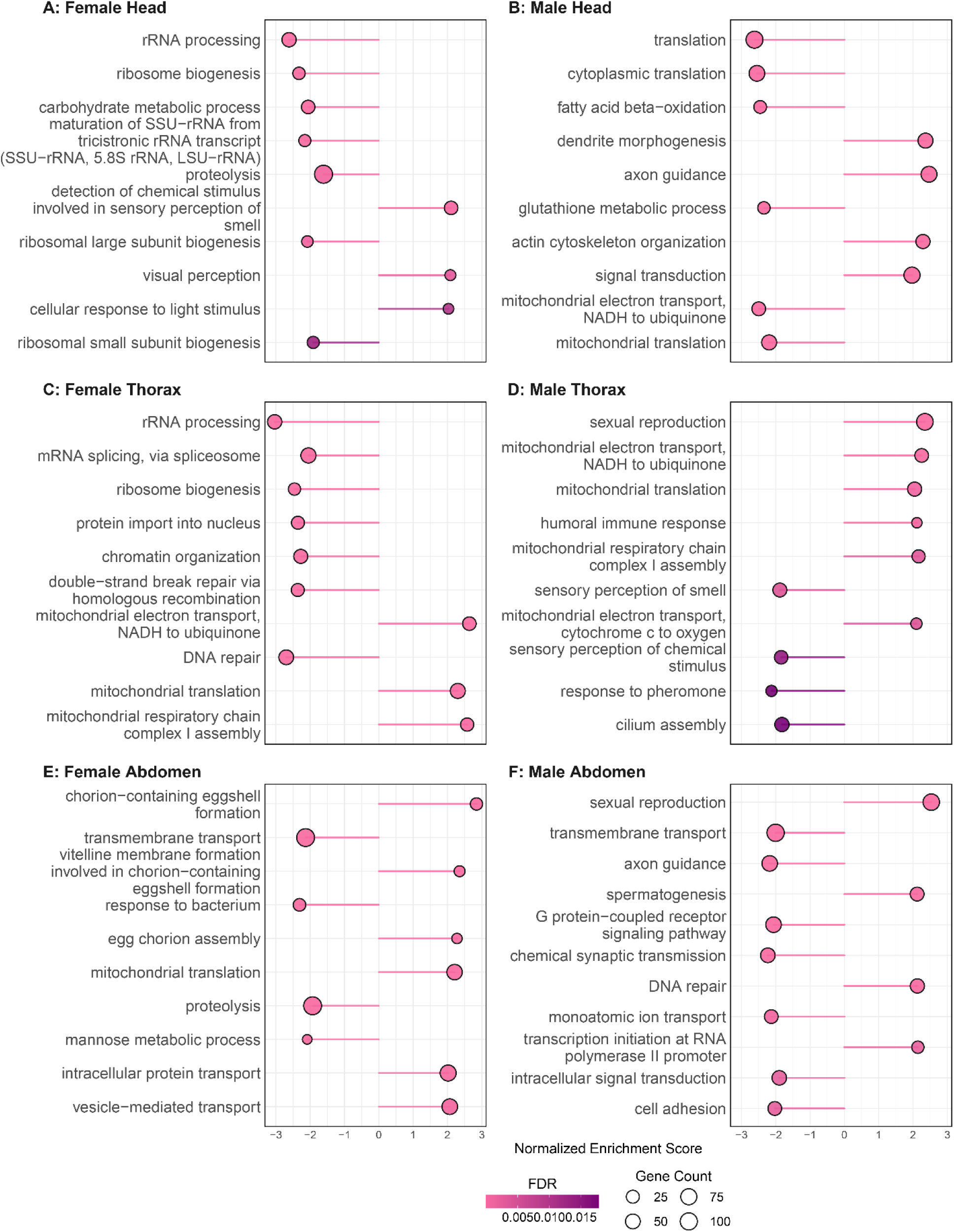
GSEA based on the Wald statistic of the interaction model. NES provides an estimate of the magnitude and direction of pathway-level differences in rapamycin response between *D. simulans* and *D. melanogaster*. As the sign of the Wald statistic reflects the direction of the additional transcriptomic effect of rapamycin in *D. simulans*, positive NES values indicate biological functions *more* activated (or *less* suppressed) by rapamycin in *D. simulans*, while negative NES values indicate functions are more strongly suppressed in *D. simulans* than in *D. melanogaster*.

In the thorax, mitochondrial-related processes were more positively affected by rapamycin in both males and females of *D. simulans* than *D. melanogaster*, contrasting with the pattern observed in male heads. Among negatively affected processes in *D. simulans* thoraces, nucleic acid pathways and ribosome biogenesis were significant in females, while immune and antibacterial defense processes were repeatedly enriched in males. Finally, the abdomen exhibited a particularly striking divergence in mTOR regulation of reproductive processes with *D. simulans* being more resistant to rapamycin-induced suppression (as indicated by positive NES values). Together, these results indicate that mTOR’s regulatory influence on core, tissue-specific pathways has diverged over the relatively short evolutionary timescale separating *D. simulans* and *D. melanogaster*.

## Discussion

Here, we used the mechanistic target of rapamycin (mTOR) pathway to study regulatory evolution under selective constraint. mTOR integrates nutrient and energy cues to coordinate growth, metabolism, and autophagy, and is highly conserved across eukaryotes, reflecting its critical role in cell physiology. By treating *D. melanogaster* and *D. simulans* with rapamycin in a fully factorial design, we quantified divergence in mTOR-mediated transcription across both sexes and three different tissues.

Our results provide clear evidence of divergence in the transcriptional effects of the mTOR signaling pathway, despite the close phylogenetic relationship between the species. We find that homologous tissues retained broadly similar expression profiles (indicated by low *d_cor_* values between transcriptomes), consistent with previous estimates (Coolon, et al. 2014) and with accumulating reports of evolutionary conservation of tissue-specific expression (Sudmant, et al. 2015). However, *D. simulans* and *D. melanogaster* exhibited clear differences in their transcriptional responses to mTOR inhibition, as indicated by high divergence values between rapamycin-induced changes in gene expression (i.e., log_2_FC). We also observed limited overlap between differentially expressed genes (Figure 3A) or enriched biological processes identified by GSEA (Figure 3B). Finally, the analysis of treatment-by-species interactions identified hundreds of genes and pathways with divergent mTOR effects. Given the close relationship between the two species, our study shows that even core signaling pathways can exhibit regulatory flexibility, making them amenable to rapid evolutionary change (Wray 2007; Fraser 2011). Furthermore, our results add to the accumulating evidence that adaptive evolution in conserved pathways often involves regulatory changes that may not directly alter essential cellular machinery (Enattah, et al. 2002; Chan, et al. 2010; Martin, et al. 2012).

Our study further reveals that the tempo of divergence in mTOR-mediated transcriptional regulation differs across tissues and between sexes. Analyses of the first-order effects of rapamycin showed that mTOR effects on gene expression are more evolutionarily labile in females than in males, and in heads than in thoraxes or abdomens. This is especially noteworthy since several earlier studies have found the rates of gene expression evolution to be slower in the nervous tissues than in other organs (Brawand, et al. 2011), generally faster in males than in females (Ranz, et al. 2003; Ellegren and Parsch 2007), and particularly rapid in male reproductive organs (Haerty, et al. 2007; Cridland, et al. 2020). However, divergence in baseline expression levels was consistent with previous findings: *d_cor_* values were generally lower in females than in males (especially in the abdomen), and the male abdomen showed the highest level of divergence among male tissues. Thus, the patterns of divergence in rapamycin response appear specific to mTOR-regulated transcription rather than reflective of genome-wide transcriptional evolution. Our results, therefore, suggest that mTOR-mediated transcriptional regulation is subject to differing selective pressures across tissues and sexes, consistent with the distinct roles mTOR plays in different cellular contexts (Laplante and Sabatini 2012; Baar, et al. 2016; Saxton and Sabatini 2017; Regan, et al. 2022; Raynes, et al. 2024; Breuillard, et al. 2025).

Pathway-level analyses of rapamycin-induced transcriptional effects provide further detail on the mode of divergence. Comparisons of enriched GO terms identified by GSEA showed that all female tissues and male heads were markedly more diverged than the male thorax and abdomen. Analyses of normalized enrichment scores (NES) confirmed that female heads and thoraxes, as well as male heads, exhibited the highest divergence, suggestive of divergent directional selection on mTOR-regulated functions. By contrast, mTOR regulation in the female abdomen, despite showing a similarly high level of divergence at the gene level, appeared more conserved at the pathway level, as evidenced by the significantly lower divergence between NES values. This pattern of high divergence at the gene level but low divergence at the pathway level may have several possible explanations. It may reflect greater robustness or redundancy (Wagner 2005) of the mTOR pathway in the female abdomen, allowing stabilizing selection to maintain function despite changes at the gene level. Another possibility is that compensatory evolution (Poon and Otto 2000) has preserved pathway functionality through offsetting alterations in different genes in the network. In any case, mTOR regulation in the female abdomen appears to evolve under a markedly different selective regime compared to other female tissues. Finally, mTOR regulation in the male thorax and abdomen appears conserved at both gene and pathway levels, as evidenced by low NES divergence. This conservation of regulatory effects suggests that the mTOR network is either under stronger evolutionary constraint and maintained by stabilizing selection, or subject to weaker directional selection than in other tissues.

Building on the analyses of first-order effects of rapamycin, we modeled species-by-treatment interactions to identify genes and pathways with species-dependent responses to mTOR inhibition. Pathway enrichment revealed that the biological processes exhibiting divergent effects of mTOR varied across tissues and sexes, consistent with tissue- and sex-specific patterns in the tempo and mode of divergence observed earlier. In the head, we found protein synthesis and carbohydrate metabolism, along with processes related to nervous system development and sensory perception to exhibit significant species-by-treatment interaction effects, indicating that mTOR-mediated transcriptional control of these critical processes has diverged between *D. simulans* and *D. melanogaster*. Interestingly, mTOR metabolic functions were less suppressed by rapamycin in *D. simulans*, while nervous-system related processes were more strongly suppressed than *D. melanogaster*. Because mTOR plays a key role in neuronal metabolism and development (Takei and Nawa 2014; Switon, et al. 2017), the divergence in mTOR signaling may reflect changes in sensory physiology or metabolic demands of the nervous system, potentially linked to ecological or behavioral differences (e.g., in host preference or courtship).

In the thorax, where females exhibited significantly faster divergence than males in earlier analysis, they also showed a greater diversity of enriched biological processes in the interaction-term GSEA. Enriched pathways in female thoraxes reflected mTOR roles in cell metabolism and growth, including rRNA processing, ribosome biogenesis, mitochondrial function, and DNA replication. In contrast, males showed very limited enrichment, sharing some mitochondrial processes with females, and only modest enrichment of immune and sensory pathways. This sex-specific pattern of divergence in mTOR-dependent regulatory effects may reflect differential constraints on muscle physiology in the two sexes.

Finally, the abdomen was characterized by relatively fast gene-level divergence in females but only moderate pathway-level divergence, and much more conserved mTOR-mediated effects in males. GSEA of the interaction term revealed that the female abdomen was enriched for reproductive, metabolic, and transmembrane transport processes, consistent with the functional roles of the ovaries, fat body, and gut. Males, despite exhibiting low divergence between species in earlier analyses, displayed broad GSEA enrichment, including neuronal signaling, immune responses, and amino acid metabolism. Notably, reproduction-related pathways were less inhibited by rapamycin in *D. simulans* than *D. melanogaster*, as indicated by positive NES values in both males and females (Figure 5). This suggests that *D. simulans* may have evolved to sustain reproduction under reduced mTOR signaling, potentially maintaining fecundity in lower-resource environments. Future studies will be needed to test this hypothesis though.

In conclusion, our results show that transcriptional regulation mediated by a highly conserved pathway like mTOR can nonetheless exhibit evolutionary divergence even between closely related species. Moreover, we find that both tempo and mode of divergence in mTOR-mediated transcription vary across tissues and sexes, suggesting distinct, context-dependent selective regimes. Enrichment analyses further reveal that species divergence in mTOR signaling has affected core mTOR-regulated pathways. The pattern of tissue- and sex-specific divergence supports the view that regulatory modularity can facilitate adaptation even in conserved pathways like mTOR (Wagner, et al. 2007; Clune, et al. 2013), allowing certain functions to diversify in specific contexts without compromising core, selectively constrained functions elsewhere.

## Materials and Methods

### Fly stocks and rapamycin treatment

We used the *D. melanogaster Oregon R* strain previously described and sequenced in (Raynes, et al. 2024). *D. simulans C167.4* (Davis, et al. 1996a) was previously described in Rand et al. (2006). All stocks were maintained at 25°C under a 12 h light-dark cycle. Standard laboratory food was prepared with 5.2% cornmeal, 2.5% yeast, 11% sugar and 0.85% agar. Tegosept (20% w/v in 95% ethanol) was added to a final concentration of 1.2% as a fungicidal preservative. Age-matched *D. melanogaster* and *D. simulans* mated adults were reared on standard food for five days, separated by sex into cohorts of 30, and transferred to vials with standard lab food or food supplemented with 200 μM rapamycin (added to the ethanol solution of Tegosept). After three days of treatment, flies were flash-frozen, dissected, placed in chilled TRIzol, and homogenized at 30 Hz for 4 min in a TissueLyser (Qiagen). Total RNA was extracted using RNeasy columns (Qiagen). RNA sequencing and initial preprocessing was performed by BGI.

### RNA-seq Data Preprocessing

Reads were aligned using STAR v2.7.10b (Dobin, et al. 2013) in 2-pass mode. In the first pass, *D. melanogaster* reads were aligned to the BDGP6.32 reference genome obtained from Ensembl, release 109 (Cunningham, et al. 2022). *D. simulans* reads were aligned to the annotated genome of the M252 strain (Palmieri, et al. 2015). In the second pass, reads were realigned to the respective reference genomes using splice junctions identified in the first pass. The *D. melanogaster* annotation (Drosophila_melanogaster.BDGP6.32.109.gtf) was obtained from Ensembl, release 109. The *D. simulans* annotation (dsim-M252-popgen-ann-r1.1.gtf) was obtained from https://doi.org/10.5061/dryad.ng95t. BAM files (sorted by coordinate) generated by STAR were indexed using Samtools v1.16.1 (Li, et al. 2009), and reads were counted using featureCounts (Liao, et al. 2014) from the Subread package v2.0.3. Only orthologous genes present in both species were retained for downstream analysis. All preprocessing was conducted using computational resources at the Center for Computation and Visualization, Brown University.

### RNA-seq Data Analysis

Read counts were imported into R v4.3.0 (Team 2023) for further analysis. One *D. melanogaster* female head control library, sequenced separately due to RNA quality issues, was identified as an outlier by MDS analysis and excluded from the study. Dispersion estimation, normalization, and differential expression testing were performed using DESeq2 v1.42.0 (Love, et al. 2014) with default parameters (lfcThreshold = 0 and alpha = 0.05). First-order effects of rapamycin were assessed using a model of the form Expression ∼ TreatmentSpecies, where TreatmentSpecies is a combination of treatment and species, followed by treatment–control contrasts for each species. Resulting *p*-values were then adjusted using independent hypothesis weighting (IHW) (Ignatiadis, et al. 2016). Genes with adjusted *p*-values < 0.05 were considered significantly differentially expressed. Second-order, interaction effects of rapamycin and species were tested using a model Expression ∼ Treatment × Species. *p*-values were again adjusted using IHW. Gene set enrichment analysis (GSEA) was performed using clusterProfiler package (Wu, et al. 2021). Genes were ranked by the Wald test statistic.

Biological process categories were considered significantly enriched at adjusted *p*-values < 0.05 using the Benjamini-Hochberg procedure.

## Data Availability

The raw RNA-seq reads generated in this study are available in the NCBI Sequence Read Archive (SRA) (https://www.ncbi.nlm.nih.gov/sra) under BioProject accession: PRJNA1107595. R scripts used in the analyses are available at https://github.com/yraynes/Sim-Mel-Divergence.

## Acknowledgements

We thank members of the Rand laboratory at Brown University for helpful discussion. This work was conducted using computational resources of the Center for Computation and Visualization, Brown University.

