## Supplementary material for "Evolutionary Divergence of mTOR-mediated Transcriptional Regulation Between *Drosophila melanogaster* and *Drosophila simulans* is modulated by sex and tissue": Supplemetal Figures

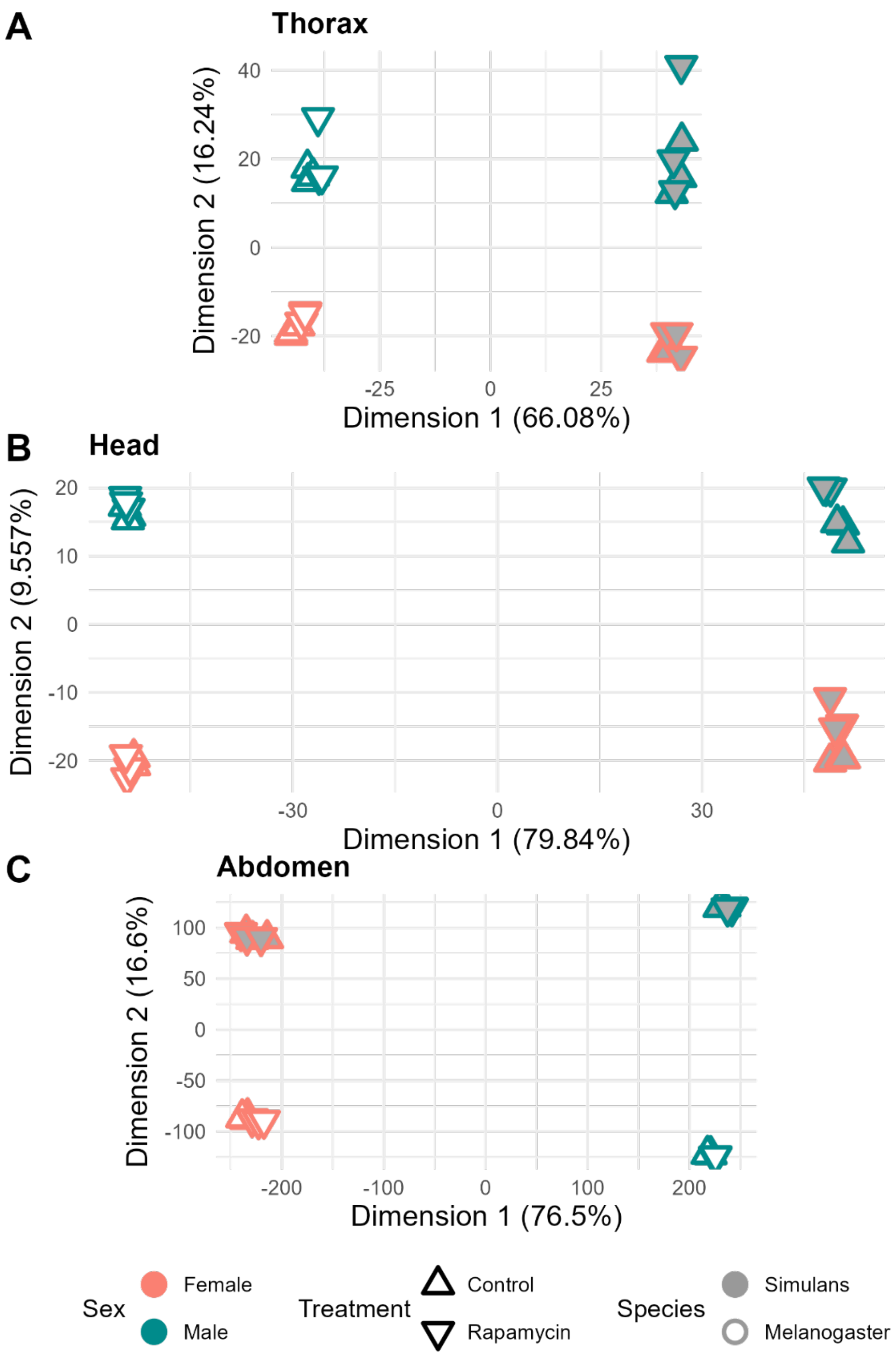


Figure S1: Multidimensional scaling (MDS) analysis of variance-stabilized transcriptome data partitioned by tissue. The relative impact of sex and species on transcription varied by body part: the difference between species explained most of the transcriptional variance in the head (PERMANOVA $R^2$ = 79.4% for species and 9.4% for sex) and thorax (PERMANOVA $R^2$=66% for species and 15.2% for sex), while the difference between sexes had the greater influence on transcription in the abdomen (PERMANOVA $R^2$=16.3% for species and 76.4% for sex).

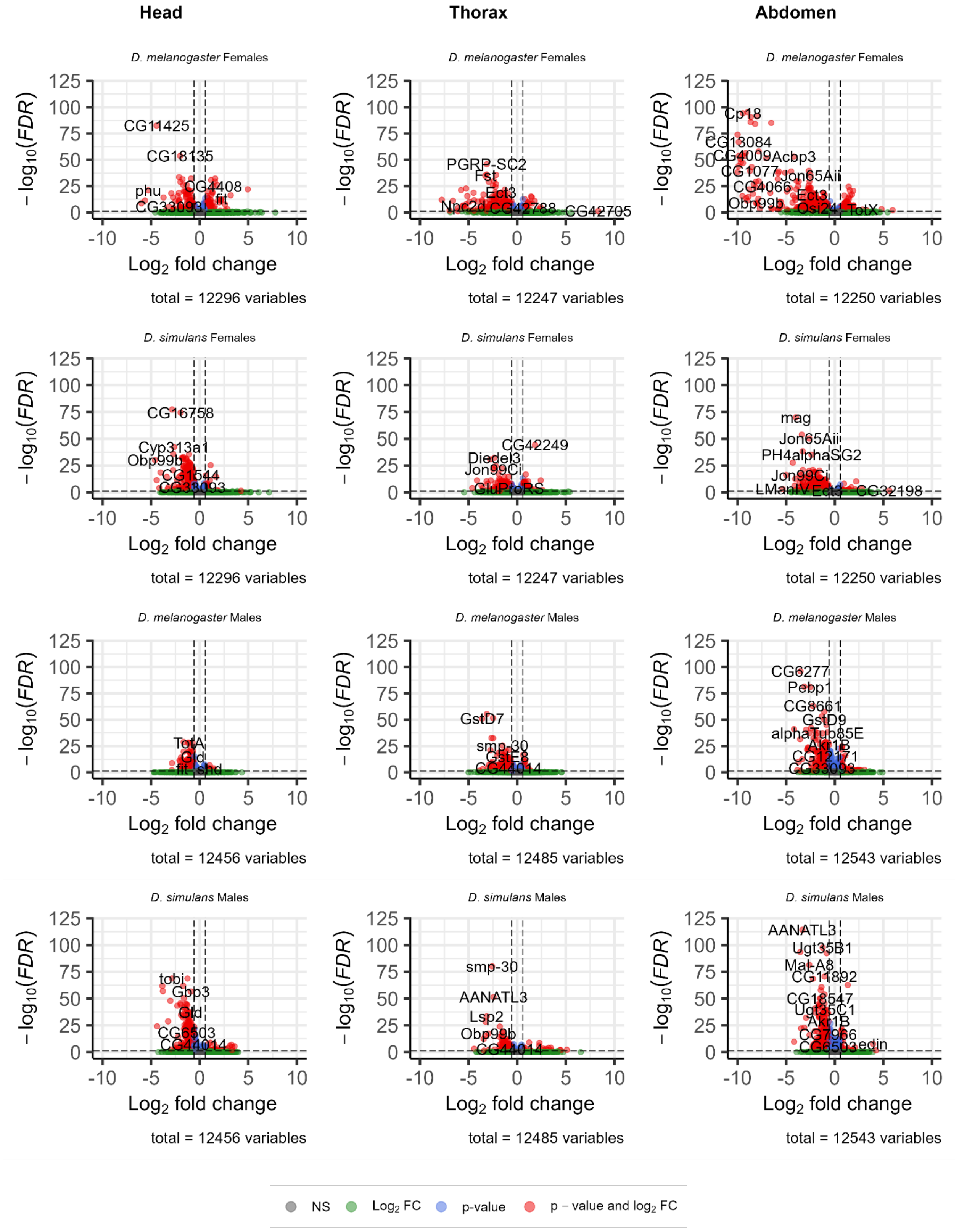


Figure S2. Rapamycin-induced transcriptional responses across tissues, sexes, and species.

Volcano plots show differential gene expression in response to rapamycin treatment for each tissue (head, thorax, abdomen), sex, and species. Genes are colored using the following categories: 1) FDR < 0.05, 2) $log_2FC > 0.58$, 3) FDR < 0.05 and $log_2FC > 0.58$, and 4) genes not significantly affected by rapamycin.

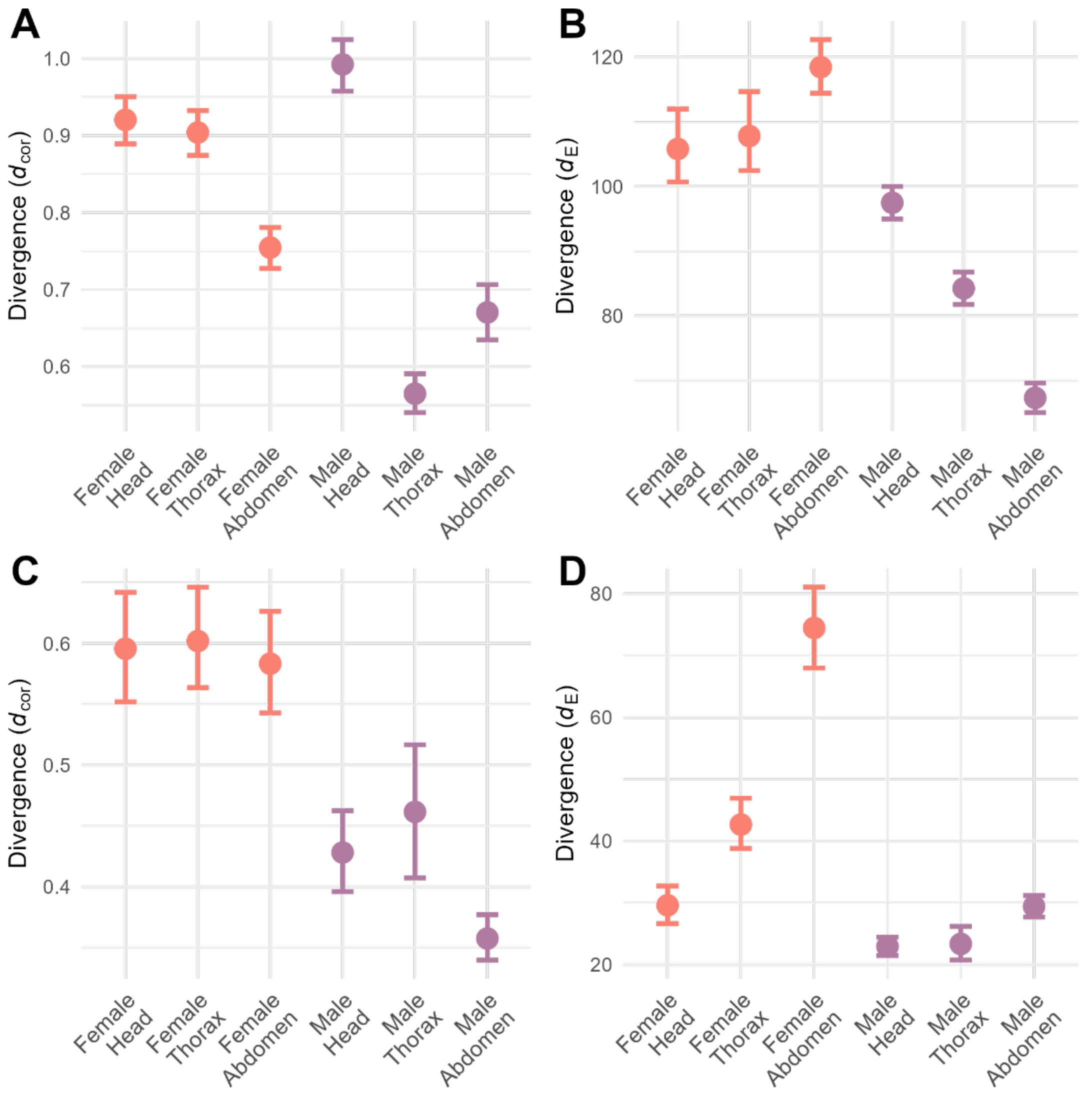


Figure S3: Gene-level divergence in rapamycin transcriptional effects: A-B) Divergence quantified using only genes with $|\log_2 FC| > 0.58$ (reflecting an approximately 1.5-fold change in expression) in at least one species. C-D) Divergence quantified using only DEGs with statistically significant $\log_2$FC (FDR < 0.05) in at least one species.

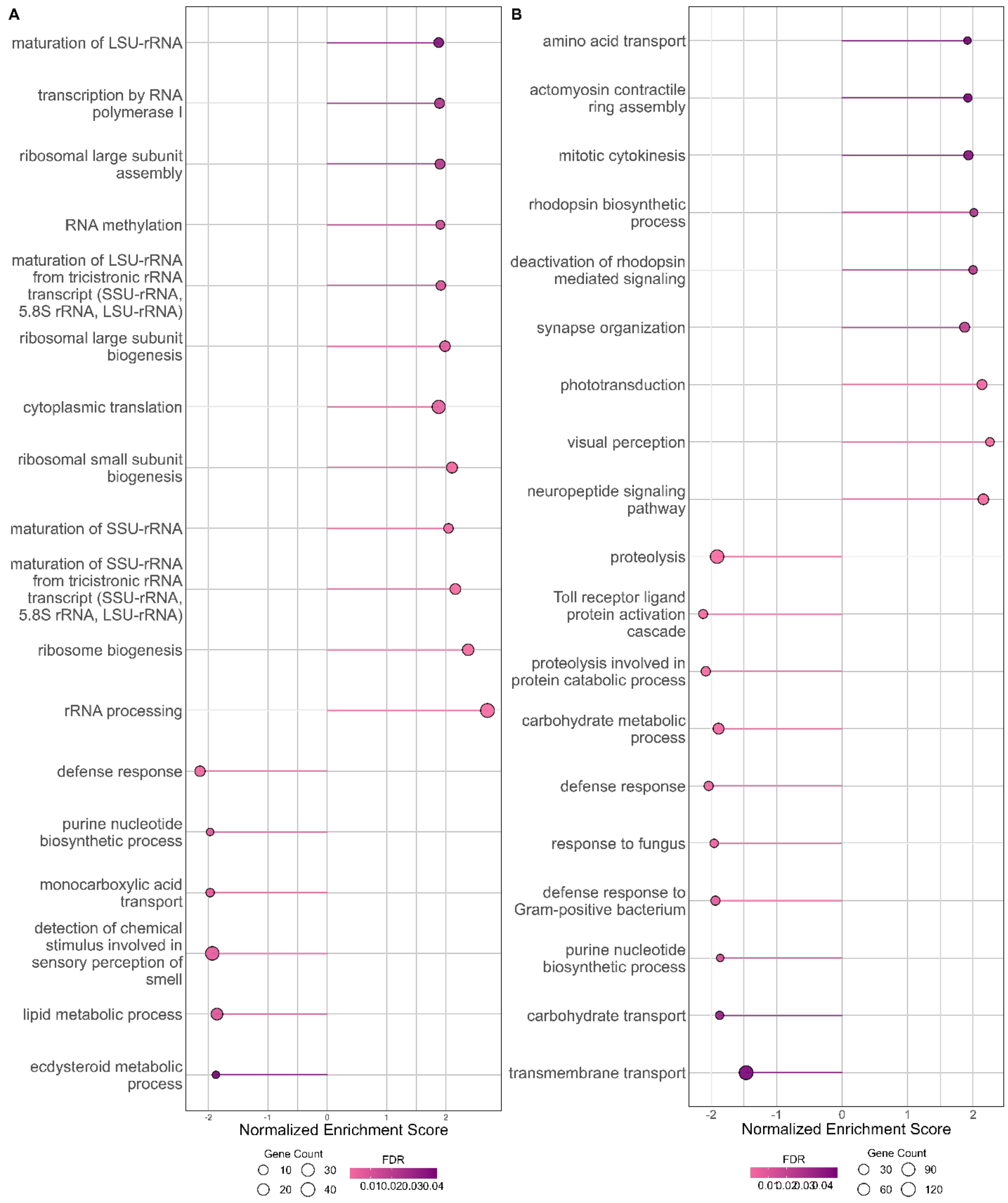


Figure S4: GSEA enrichment results in female heads: A) *D. melanogaster* B) *D. simulans*. All significant GO terms shown (FDR < 0.05)

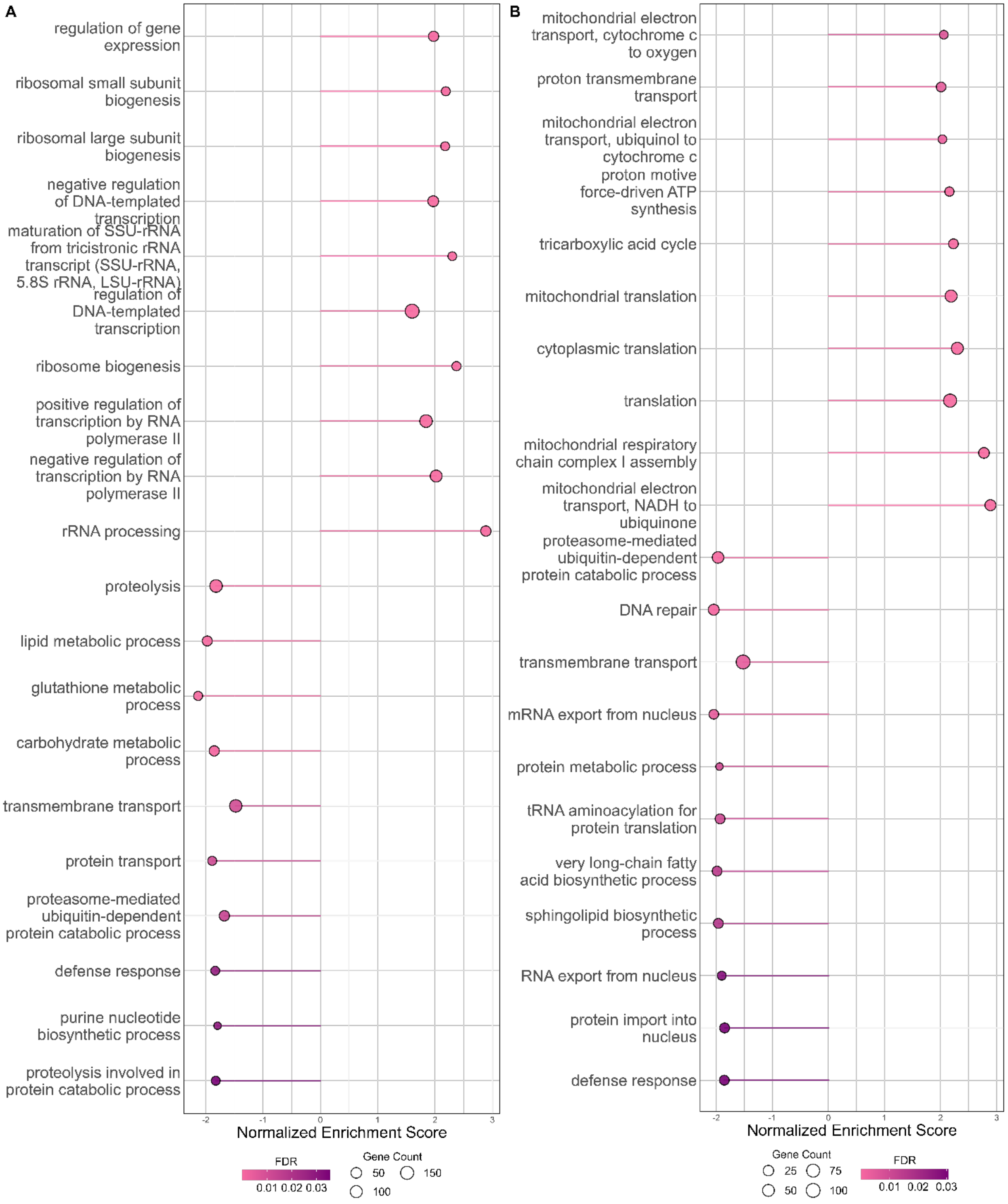


Figure S5: GSEA enrichment results in female thoraxes: A) *D. melanogaster* B) *D. simulans*. The top ten GO terms most significantly suppressed or activated by rapamycin, ranked by lowest FDR, are shown.

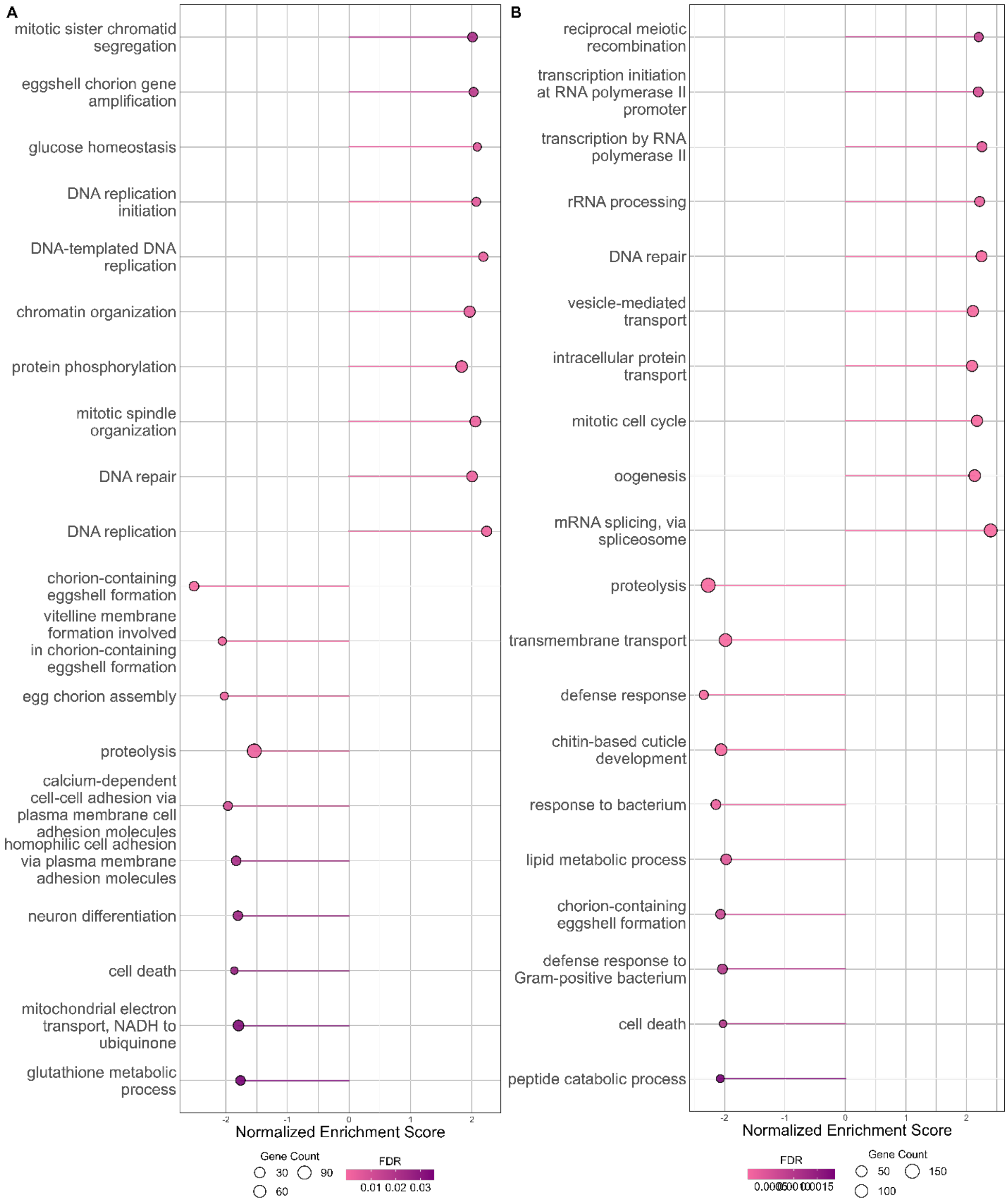


Figure S6: GSEA enrichment results in female abdomens: A) *D. melanogaster* B) *D. simulans*. The top ten GO terms most significantly suppressed or activated by rapamycin, ranked by lowest FDR, are shown.

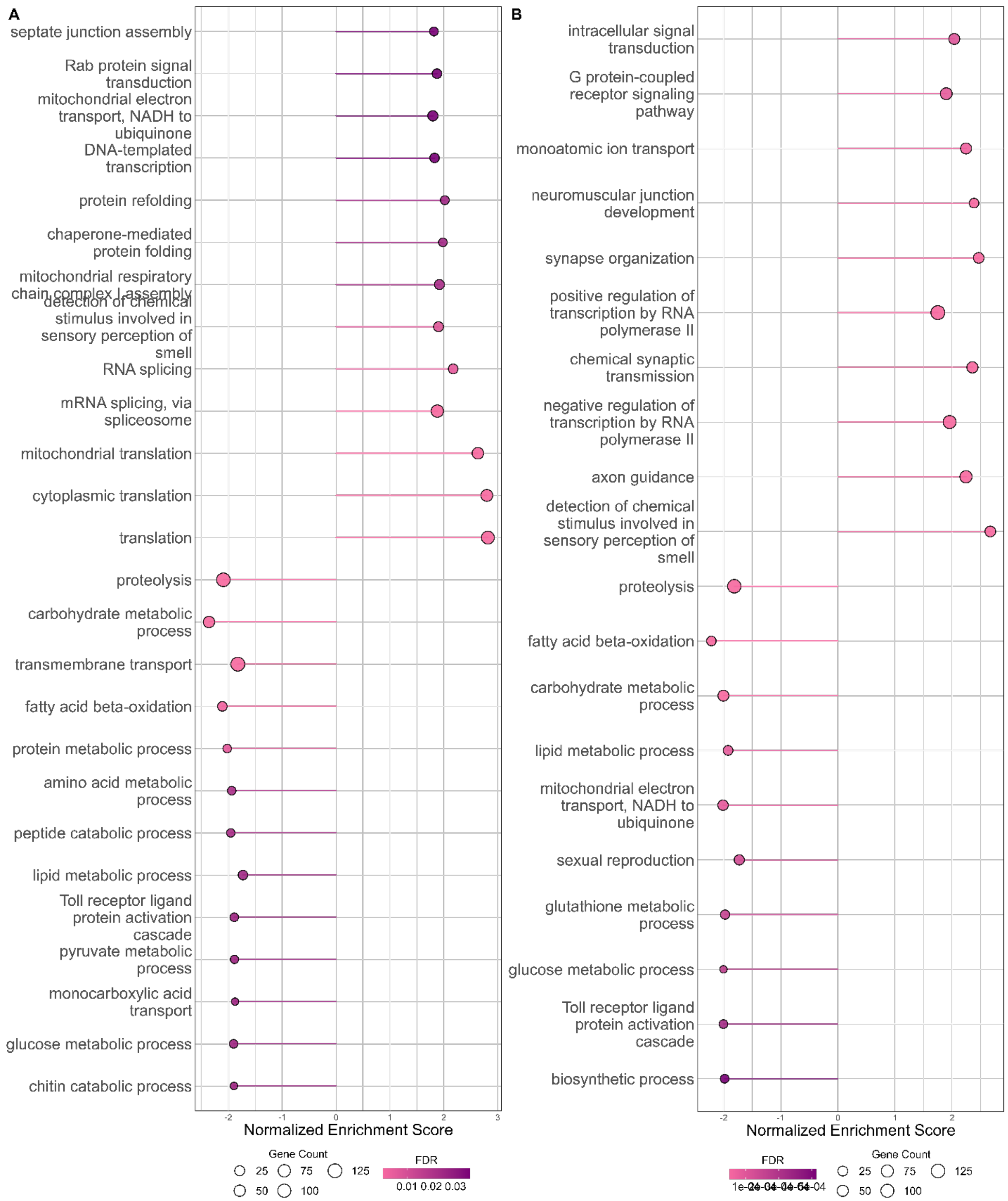


Figure S7: GSEA enrichment results in male heads: A) *D. melanogaster* B) *D. simulans*. The top ten GO terms most significantly suppressed or activated by rapamycin, ranked by lowest FDR, are shown.

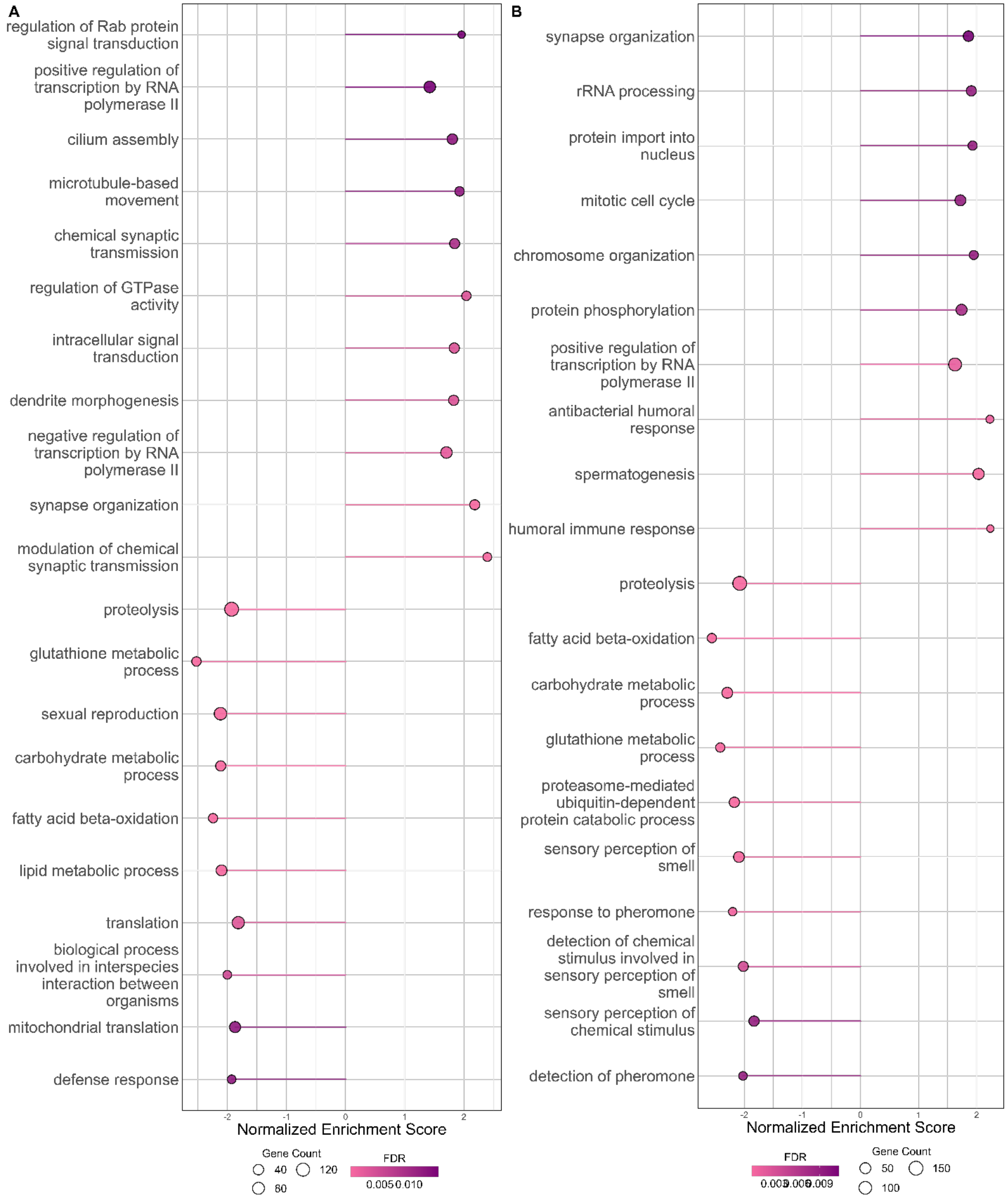


Figure S8: GSEA enrichment results in male thoraxes: A) *D. melanogaster* B) *D. simulans*. The top ten GO terms most significantly suppressed or activated by rapamycin, ranked by lowest FDR, are shown.

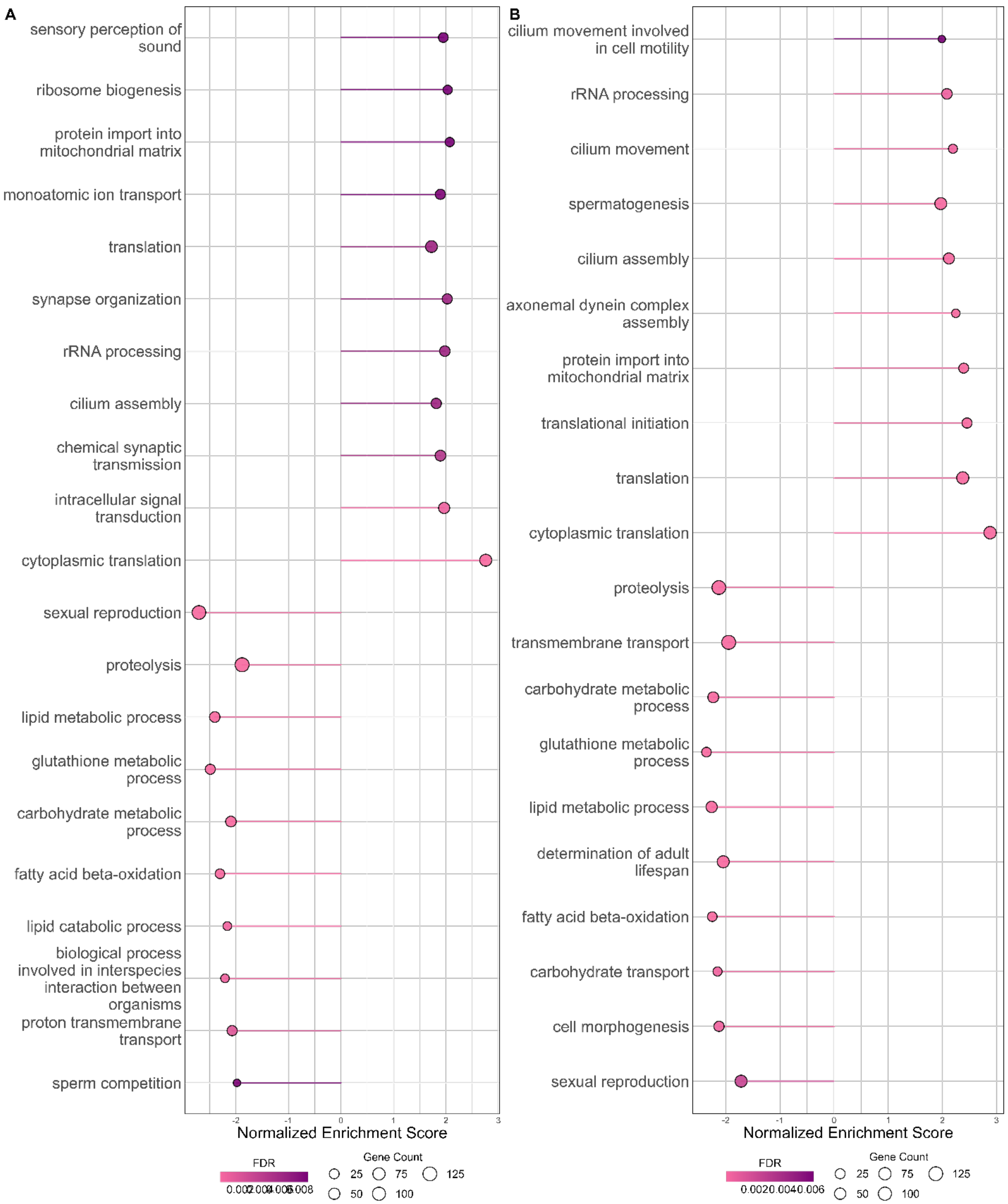


Figure S9: GSEA enrichment results in male abdomen: A) *D. melanogaster* B) *D. simulans*. The top ten GO terms most significantly suppressed or activated by rapamycin, ranked by lowest FDR, are shown.

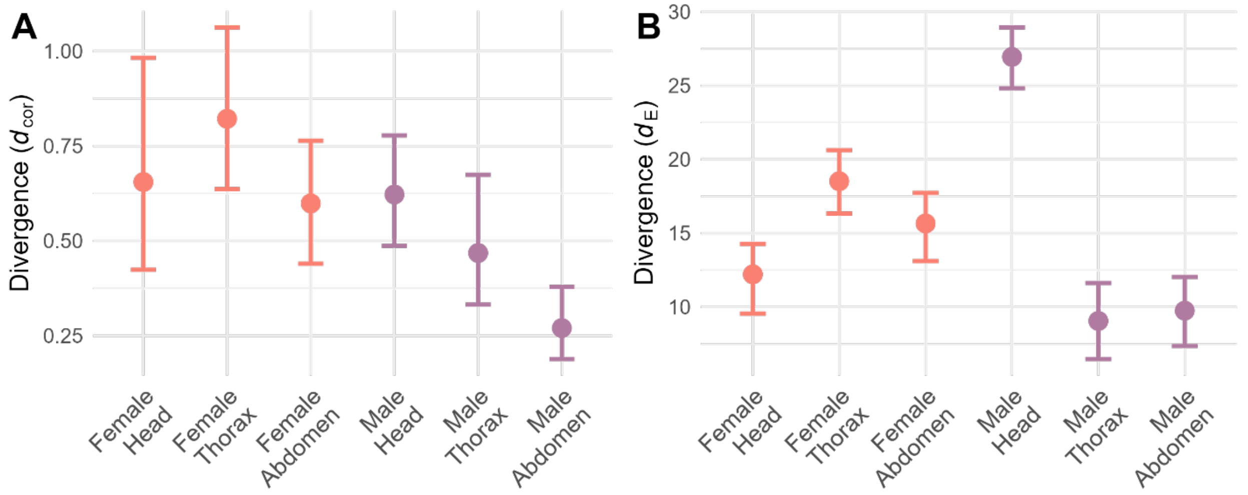


Figure S10: Divergence in rapamycin response at the pathway level. A) $d_{cor}$ $(1 - \rho)$ between *D. melanogaster* and *D. simulans* NES values. B) $d_E$ (Euclidean distance) between *D. melanogaster* and *D. simulans* NES values. All calculated using only the pathways significant in either species. Error bars indicate 95% bootstrap confidence intervals (10,000 replicates with replacement).

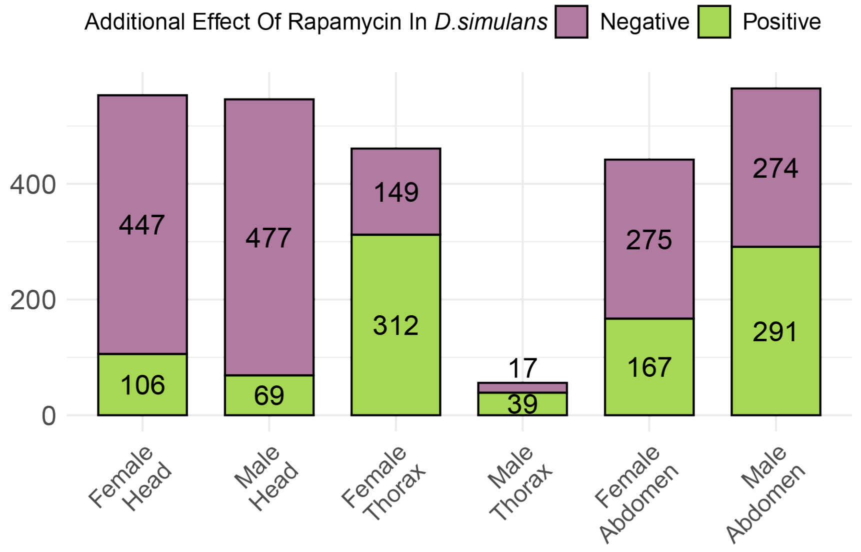


Figure S11: DEGs with significant treatment-by-species interaction terms, classified by the sign of the *additional* effect of rapamycin in *D. simulans* relative to *D. melanogaster* (represented by $\log_2$FC for the interaction term): positive $\log_2$FC indicates that rapamycin increases expression more (or suppresses it less) in *D. simulans* than in *D. melanogaster* and a negative $\log_2$FC suggests that rapamycin increases expression less (or suppresses it more) in *D. simulans*. It is calculated as $\log_2$([*D. simulans* expression in Control]/[*D. simulans* expression in Rapamycin]) - $\log_2$ ([*D. melanogaster* expression in Control]/[*D. melanogaster* expression in Rapamycin]). The number of DEGs exhibiting significant treatment-by-species interactions was broadly similar across most tissues (except in the male thorax). However, the direction of interaction effects varied between tissues. In both male and female heads, most significant DEGs were more negatively affected by rapamycin in *D. simulans* than in *D. melanogaster*. In contrast, the distribution of interaction effects was more balanced in the abdomen, suggesting that the effects of mTOR on expression have changed in both directions: stronger in *D. simulans* for some genes and in *D. melanogaster* for others. The female thorax was skewed towards DEGs activated more (or less suppressed) in *D. simulans*, while the male thorax contained very few interaction DEGs, consistent with a more conserved response to mTOR inhibition.
